# Seasonal differences in predation risk among seagrass epifauna species stabilize community-level predation over time

**DOI:** 10.64898/2026.01.05.697804

**Authors:** Claire E. Murphy, John J. Stachowicz

**Affiliations:** Department of Evolution and Ecology, University of California, Davis, California, USA; Bodega Marine Laboratory, University of California Davis, Bodega Bay, California, USA

**Keywords:** predator-prey interactions, seasonal variation, seagrass, epifauna

## Abstract

Predation risk varies through space and time due to changing refuge quality, predator communities, and prey traits. Despite this, ecological research is often focused on measuring average predation risk at the community level. While this can give important information about overall trophic transfer and ecological efficiency, it ignores differences in predation risk among prey species within a community, which may be important determinants of species coexistence and local diversity. We used crustaceans associated with temperate seagrass in Northern California to explore the relationship between seasonal variation in among-species and community-level predation risk for a community of morphologically distinct prey. We measured predation risk of the four most abundant and widespread prey species at six field sites every two to six weeks for one year. At the community level, sites differed significantly in their annual variation in predation risk, and these differences were correlated with the amount of variation in the among-species predation risk. When there was more within-year variation in predation risk among the four prey species, predation risk at the community level was more stable across the year. On the other hand, when each prey species in the community had similar levels of predation risk throughout the year, predation as a community-level process was much more seasonal and variable. Variation in predation risk also changed across a gradient of seagrass cover, a proxy for refuge quality. Sites with greater seagrass cover had less annual variation in community-level predation risk and more variation in predation risk among the four species at any given time point. In contrast, at sites with less eelgrass, all species were consumed at the same rate throughout the year, suggesting previously demonstrated differences in antipredator strategies among species are less relevant in the absence of habitat-forming species. We suggest that larger species-specific differences in predation risk throughout a year result in a more stable level of predation risk for the whole community, and that this may be driven by increased refuge provided by seagrass habitat mediating different prey species’ relative levels of susceptibility to predation.

## Introduction

The strength of predation, and thus the risk that a prey community experiences, is dependent on three main factors: predator communities (composition and abundance), the refuge available in the environment (type and amount), and prey traits. The same prey community can experience different levels of risk across geographic gradients as refuge availability and predator communities shift, such as through changing habitat structure (e.g., Schneider 2001, Warfe and Barmuta 2004), predator composition and behavior (e.g., Brown-Saracino et al. 2007), or even hunting regulations (e.g., Selden et al. 2017). On top of this, within a prey community different species often have unique traits or behavioral responses to these changing environmental conditions, and a context that favors the antipredator responses of one species may make another more vulnerable (Ives and Dobson 1987). Furthermore, habitat or predator community contexts are not static – seasonal shifts can change which antipredator responses are most effective and as a result, which prey species experience the most risk (Gratton and Denno 2003, Mappes et al. 2014).

Despite this potential for large variation in risk, understanding the aggregate “community-level predation risk” (i.e., the average risk each member of the prey community experiences, regardless of species identity; the solid lines on Figure 1), can give important information about trophic transfer and ecological efficiency. Across a year, community-level predation risk may fluctuate in seasonal environments as predator communities or refuge availability change (Figure 1A). In contrast, more stable environments have more consistent predator communities and refuge availability and so prey communities might experience constant levels of predation risk (Figure 1B & 1C). However, consistency in community-level predation risk throughout a year could arise from two very different mechanisms (Figure 1B & 1C). In one scenario, there are equally consistent, hierarchical predation patterns for each individual species in the community (Figure 1B). Alternatively, low seasonal variation in community-level predation can also be the result of compensatory dynamics in which the risk hierarchy among species varies considerably across the year, but the averaged total risk is relatively constant (Figure 1C).

**Figure 1.**
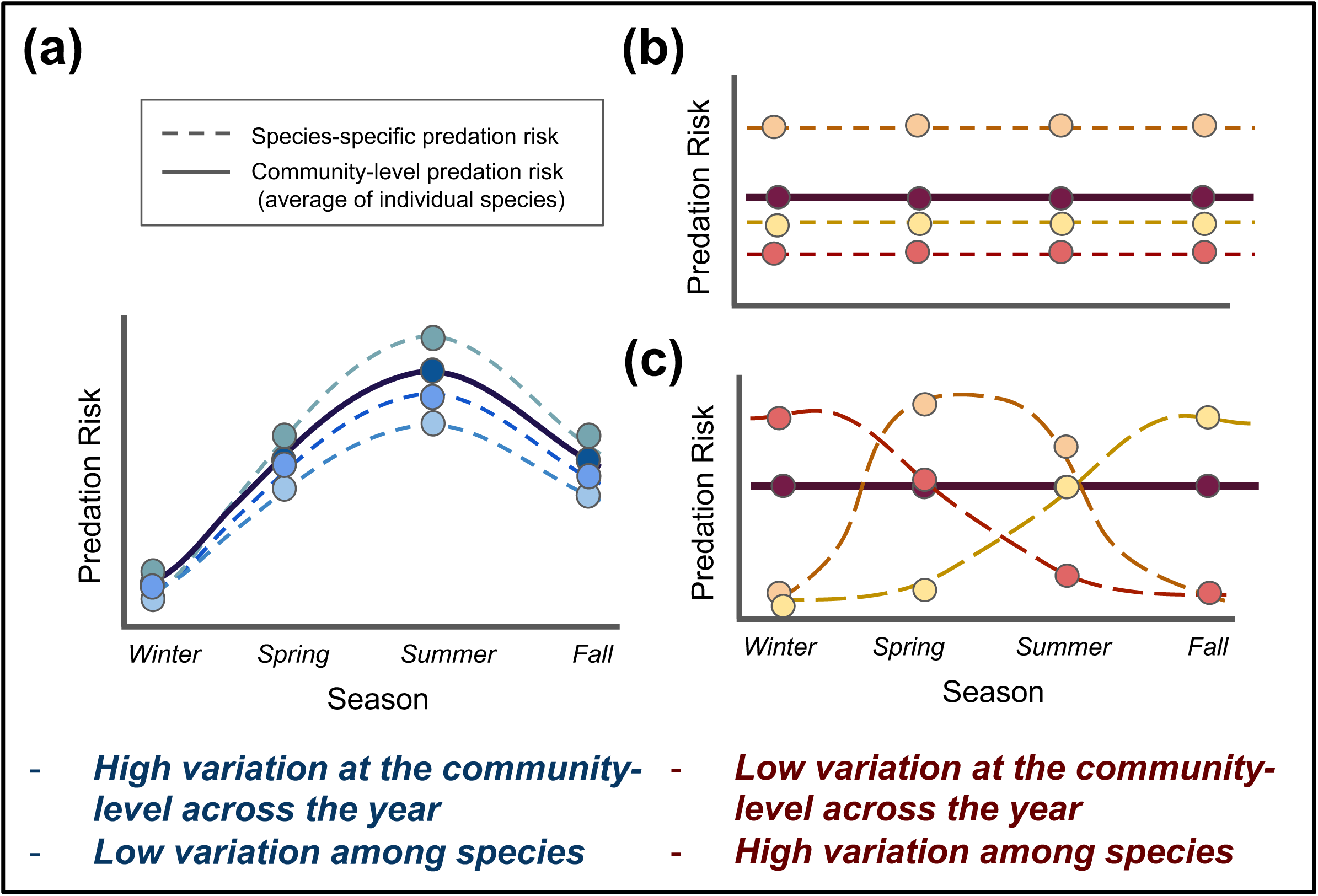
Hypothetical scenarios demonstrating the two extremes of variation in community-level predation risk across a year and the individual species-level patterns that can lead to them. Each different colored line represents a different species and the predation risk it experiences across a year. The solid lines represent the community-level predation risk, or the average risk all species experience at any given time point. Panel **(a)** represents a prey community that experiences high annual variation in community-level predation risk, while panels **(b)** and **(c)** represent two different scenarios that lead to stable community-level predation risk across a year.

This variation in species-specific predation risk across a prey community could play an important role in prey co-occurrence patterns and how they change across seasons or regions. At a site with a constant hierarchy in predation risk among species, the most vulnerable prey might be unable to persist, having to rely on high levels of dispersal or reproduction to counteract predation (Amezcua and Holyoak 2000). Alternatively, if prey experience complementary and seasonally variable amounts of risk, it may be that no one species is able to remain dominant due to regular periods of high predation, promoting prey diversity and stability (Rakowski et al. 2021). Despite these differences, both scenarios would display stable community-level predation risk across the year. Furthermore, because species differ in predator-avoidance strategies, a generic prey item may misrepresent not only predation risk to individual species and but also the total community-level risk, if used as a proxy (Mihalitsis et al. 2021). Thus, while aggregate risk is a useful measure of total trophic transfer at a site, understanding predation’s effects on the whole community requires separate assessments of risk for each prey species. These species-specific predation risk data are also critical to connect patterns of variation in predation risk to prey population and community dynamics. Despite this, high-frequency data measuring the relative predation risk among dominant species are rarely available.

Temperate seagrass meadows systems, such as those dominated by eelgrass (*Zostera marina*), support invertebrates with a diversity of antipredator adaptations that may be effective against different predators (Lürig et al. 2016). These eelgrass beds are known to vary spatially and seasonally in invertebrate community composition (Ha and Williams 2018, Gross et al. 2022), abundance and diversity of predators (Douglass et al. 2010, Ruesink et al. 2019), and habitat structure (Hovel et al. 2002, Duffy et al. 2022) and provide a tractable system in which to understand the drivers of temporal variation in predation risk in the field. Considerable research has considered how habitat amount and structure, the main form of refuge, affects the abundance and predation risk of these epifaunal invertebrates (Boström et al. 2006, Yarnall et al. 2022).

Generally, increased habitat, either measured through increased total seagrass biomass (Virnstein et al. 1984), total shoot surface area (Sirota and Hovel 2006), or percent cover (Reed and Hovel 2006), is positively associated with increased diversity and abundance of epifauna. A common explanation invoked for this is that more complex habitats result in lower predation (Sredl and Collins 1992, Hovel et al. 2021, Gigliotti et al. 2021). However, reduced community-level predation does not necessarily lead to higher prey diversity (Finke and Denno 2005, Ryberg et al. 2012), and a better understanding of how predation risk varies among species in changing habitat contexts through space and time is needed to link predation and local community composition. Additionally, the interpretation of geographic comparisons in predation risk using standard prey items (Reynolds et al. 2018, Whalen et al. 2020) requires a better understanding of the consistency of among-species variation in predation.

Here we measure predation in the field on four species of peracarid crustacean at six sites at ∼3-week intervals over the course of an entire year. We evaluate the relationships between within-year variation in predation risk and habitat amount by addressing two questions: 1) what is the relationship between seasonal variation in species-specific and community-level predation risk? and 2) how does this relationship change across a gradient of habitat amount?

## Methods

### Study sites and focal species

We conducted all experiments and surveys in the seagrass beds in Bodega Harbor and Tomales Bay, California. Both estuaries support extensive beds of *Zostera marina* (eelgrass), a widespread temperate foundation species, exhibit strong seasonal and spatial environmental changes, and host a diversity of invertebrates and fishes (Olyarnik and Stachowicz 2012, Best and Stachowicz 2014, DuBois et al. 2022, Gross and Stachowicz 2024).

We focused on four abundant, widespread, and functionally distinct species of epifauna – the caprellid amphipod *Caprella californica*, the ampithoid amphipods *Ampithoe lacertosa* and *Ampithoe valida*, and the isopod *Pentidotea resecata* that represent the wide range of predation avoidance traits found in epifaunal communities (Best and Stachowicz 2012). The caprellid amphipod, *C. californica*, occurs at high densities (Ha and Williams 2018) and reproduces quickly with large and frequent broods (Keith 1971). They are thin and inconspicuous, allowing them to blend in with the seagrass, and individually they have low caloric value (Caine 1991).

They are relatively easy for predators to capture and seem to counteract the impacts of predation through rapid reproduction and large populations (Caine 1989, Hosono 2014). In contrast, the isopod *P. resecata* grows to be one of the largest members of the grazer community (Lürig et al. 2016). They camouflage in the eelgrass by matching the blades in shape and color (Lee and Gilchrist 1972). This unique morphology may allow them to evade predation, especially when there is a high density of eelgrass blades to hide among (Best and Stachowicz 2012). Finally, the ampithoids *A. valida* and *A. lacertosa* have specialized silk glands which allow them to create temporary protective tubes within the eelgrass or macroalgae, decreasing their risk of predation (Nelson 1979). Beyond this they are neither morphologically defended nor particularly cryptic on eelgrass. All these species are generalist herbivores that consume both live and dead eelgrass, epiphytes, and macroalgae to varying degrees (Best and Stachowicz 2012).

### Predation assays

To measure predation risk, we tethered live individuals of each of the species by gluing them (using Loctite Ultra Gel Control cyanoacrylate super glue) to a 10 cm long monofilament line attached to the top of a 35 cm long clear acrylic rod, which we then embedded in the sediment, leaving about 15 cm above ground. We did this at 6 sites every 2-6 weeks (a total of 16 times from April 2022 to March 2023). At low tide at each site, we placed 20 individuals of each species along a set transect, with each tether 0.5 m apart from the next, alternating between the focal species. At the three Tomales Bay sites we included all four focal species (resulting in 40 m transects) but excluded *A. valida* in Bodega Harbor (resulting in 30 m transects) since it does not regularly occur there and may be non-native (Harper et al. 2022). We collected individuals to tether from all six sites, depending on where each species was most common and easy to collect at each time point, and occasionally supplemented this with laboratory cultures as needed. In a few instances, weather or low availability prevented the inclusion of particular sites or species at a time point.

We retrieved the tethers at low tide approximately twenty-four hours later. For each tether that was recovered, we recorded whether the prey item was missing, which we assumed meant it had been consumed. Underwater videos showed fish eating tethered individuals, demonstrating that predation was a reason for their removal (Appendix S1: Figure S1A; Video S1). Sometimes tethered individuals were crushed or ripped in half, presumably by a predator consuming them (Appendix S1: Figures S1B & S1C). In addition, in laboratory pilot experiments, no individuals of any of the four species came unglued from tethers, even after 5 days, suggesting that individuals missing from tethers in the field most likely not the result of glue failure. We discuss additional evidence in the Results that suggests that temperature or water motion were unlikely to cause the patterns we observe due to a lack of correlation between frequencies of missing prey and any environmental driver.

We calculated predation risk for each species at each site x time point combination as the proportion of tethers with missing prey items. While tethering experiments do not give absolute measures of predation risk, we can use data collected from them to understand the relative risk different species might experience, and how this risk might change through space and time (Peterson and Black 1994, Aronson and Heck 1995, Baker and Waltham 2020, Rhoades et al. 2024). Within one deployment, we inferred that the tethered species that were consumed the most at a site also had the highest risk of being consumed in the field at that point in time. Across deployments, if a particular species was missing from more tethers at one point in time relative to another, we concluded there was a greater risk of predation at that time. We acknowledge that tethering can differentially impact species and inflate the predation risks of prey with more behavioral predator avoidance strategies. Yet we observed tethered tube builders (*Ampithoe* spp.) constructing tubes in a similar manner to free untethered ones (Appendix S1: Figure S1D & S1E), and we regularly observed tethered *P. resecata* individuals clinging to blades of eelgrass to camouflage (Appendix S1: Figure S1F & S1G). Furthermore, there were points in the year where each species had very low levels of predation risk (e.g., an average proportion of 0.148 of all tethers at a site consumed at one deployment), and these did not always coincide among species (see Results). Together, this suggests that tethering did not completely eliminate prey defenses and that our data reflect relative risk of predation among species and times for the taxa and sites we used.

### Habitat surveys

Preliminary examination of predation risk data indicated that habitat amount (the percent cover and shoot density of eelgrass) one month after concluding the tethering experiments (March 2023) was associated with differences in predation risk among sites, prompting us to include measures of habitat in our analysis. However, due to the frequency and intensity of the sampling required to measure predation risk during the year of tethering, and the limited duration of low tides, we could not also feasibly measure habitat concurrently. We returned to each site every four to six weeks (n = 11) over the next year from April 2023 to March 2024. Along the same transects that we had deployed the tethers, we estimated the percent cover and shoot density of the seagrass in five 0.25 m^2^ quadrats, evenly spaced along the length of the transect.

We also noted the percent cover of any additional macroalgae or sessile invertebrates present within the quadrat. To measure percent cover, we estimated the amount of ground covered by each substrate in the quadrat, both visually and by touch. Total measures of percent cover could exceed 100% within a given quadrat if different substrates were layered on top of each other.

When multiple observers were involved, measurements were standardized before collecting data to ensure consistency in the percent cover estimates. We measured both percent cover and shoot density of the seagrass as they provide slightly different but complimentary information about the bed. Shoot density gives a measure of how complex the habitat is, while percent cover better represents the total amount of habitat. For example, the same number of shoots may yield different percent cover estimates if they are seedlings in the spring versus large adult shoots in the late summer.

We believe that these data are representative of the rank-order differences in habitat among sites during the year we conducted the tethering for several reasons. Our initial habitat survey occurred just one month after our last predation survey, and the rank order of the sites in terms of percent cover and shoot density measurements from this time point are strongly correlated with the rank order of sites based on our yearlong averages (shoot density: rho = 0.968, df = 4, p = 0.002; percent cover: rho = 0.904, df = 4, p = 0.013). Slight differences in the timing of seasonal growth and senescence of the eelgrass among sites does result in changes in the rank order of sites with respect to eelgrass cover, especially throughout the winter. To account for this, we only use these data to assess the influence of mean eelgrass cover differences among sites, rather than fine scale differences in the precise timing of peak habitat, which we do expect could vary among years. Finally, for periods where we do have multiple years of habitat cover data from the same sites (from April – July 2023 and 2024), there was a strong correlation in the rank orders of sites calculated using both the shoot densities and percent cover of seagrass between the two years (percent cover: all rho values between 0.812 and 0.943; all p values between 0.005 and 0.049; shoot density: all rho values between 0.886 and 0.943; all p values between 0.005 and 0.019). Thus, we feel comfortable concluding that the relative amount of habitat present at each of our sites was consistent during the years of our study.

In our analysis, we assessed Pearson correlations between different metrics of predation risk (see below) with the average shoot density, and the average percent cover of seagrass, bare ground, and macroalgae. We focused on spatial differences in habitat amount across the whole year, rather than seasonal variation within the year as, with respect to habitat, the sites differed to a greater degree among than within sites across the year (Appendix S1: Figure S2). Since there were large differences in the average amount of habitat between the sites, it was impossible to disentangle whether differences in the annual variation of habitat amount were driven by real differences in the level of seasonality at a site, or just by these differences in the means (which were positively correlated with variance). Habitat metrics were not confounded with site temperature, as neither the rank order of average shoot density nor average seagrass percent cover correlated with summer temperature at each site (shoot density: rho = −0.143, df = 4, p = 0.803; percent cover: rho = −0.086, df = 4, p = 0.919; temperature data from Schiebelhut et al. 2023).

### Predation and habitat analysis

We calculated seasonal variation in among-species differences in predation risk and overall aggregate community-level predation risk across a year and then assessed the relationships between these two metrics to each other and between each of them and eelgrass cover.

First, to assess whether the species all followed similar temporal patterns of predation risk (as in Figures 1A & 1B), or if the species differed in their patterns (as in Figure 1C) we compared the distributional shapes of the annual predation patterns. To do this we used a method originally designed to compare the shapes and distributions of flowering schedules of individual plants (Austen et al. 2014). This gave us a metric of how variable the seasonal predation risk was among the species in the prey community at each site.

To do this we first created a distance matrix of Kolmogorov-Smirnov (KS) test statistics. KS tests evaluate the likelihood that one distribution was sampled from another reference probability distribution. We ran pairwise KS tests to compare the distributions of predation risk through time between each species at each site (i.e. within each box in Figure 2A we analyzed how different the overall shape of each colored curve was from the others). Lower KS test statistic values indicated that the distribution of predation risk through time was more similar for two unique species x site combinations. Values can range from 0 to 1, with a value of 0 indicating that two distributions are identical. KS test statistics are sensitive to both differences in the magnitude as well as the timing of peaks between different distributions. To visualize these multivariate differences, we ran a principal coordinates analysis (PCoA) on the KS distance matrix. These KS distances do not imply a specific temporal structure or allow for conclusions about synchrony; for example, a shift in the peak of the distribution by 1 month will produce an identical KS distance regardless of whether the peaks occur in March and April or August and September. Nevertheless, they do allow clear conclusions about the extent to which the timing and magnitude of predation peaks are similar for each pair of distributions compared. We ensured that our KS distances fulfilled the triangle inequality and only interpreted our results based on the positive eigenvalues when evaluating the PCoA.

**Figure 2.**
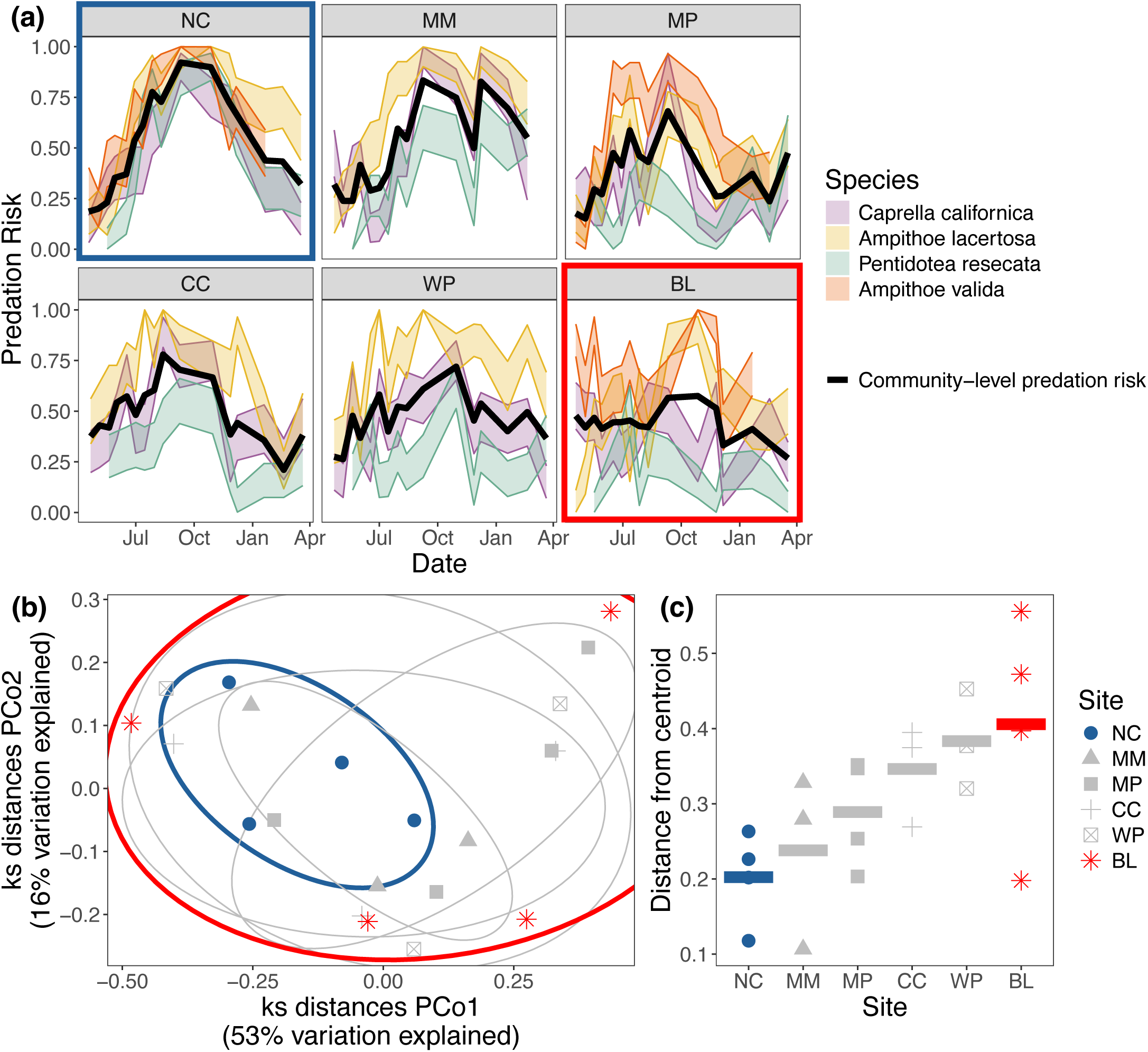
Panel **(a)** shows the seasonal patterns of predation risk for the four species community at each of the six sites. The width of the shaded lines represents the standard error around the average predation value (the average proportion of recovered tethers that had been eaten) at each time point. Each colored line represents a species, and the solid black line represents the community-level average predation risk. Sites are ordered from the site with least habitat to the site with the most. In panel **(b),** each species by site predation pattern is plotted in Kolmogorov-Smirnov (KS) multivariate space. Each point represents a distribution of predation risk across the year (e.g., one of the colored lines in **a**) for each species at every site. The distance between two points correlates to how similar those two distributions are (e.g., the similarity of the shapes of two different colored lines in **a**). Sites varied in the level of similarity of the predation patterns for each species **(c)**. In **(c)**, sites are ordered on the x-axis from the lowest amount of seagrass cover to the highest amount of seagrass cover, and the y-axis represents how spread out the predation patterns of each species at that site are in KS space (represented in **b**). Sites with more seagrass have more variation in the distributions and shape of the seasonal predation risk for each species (as in Figure 1C). The sites with the most and least dispersion in KS space are colored red and blue respectively in all three panels.

We also tested whether there were differences in the dispersion of KS distances in multivariate space, either by site or species groups, to get a measure of how variable or consistent different groups were in their distributions of predation risk across the year. We used the *betadisper* function in the vegan package (Oksanen et al. 2017) to calculate the centroid of each species or site grouping in multivariate space, based on their pairwise KS distances, and then to calculate how far the seasonal pattern of predation risk for each unique species and site combination was from those centroids (i.e., dispersion). A lower level of dispersion at a site would indicate that the predation patterns of each species may resemble Figure 1A or 1B, while larger differences would suggest the patterns are more like Figure 1C. We used the level of dispersion between species-specific predation patterns as our measure of “among-species variation in predation risk” for each site.

Next, we wanted to understand how predation risk varied across the year at the aggregate community level regardless of species-specific risk. To calculate this community-level predation risk for each site, we calculated the proportion of all tethered animals consumed at each time point, regardless of which species they were. Then, to quantify how seasonally variable predation was at the community level for each site (“community-level variation in predation risk”), we took the standard deviation of this average community predation risk across all the time points within a site. This gave us one value per site.

To assess the relationship between the annual variation in species- and community-level predation risk within a site, we calculated Pearson r correlation coefficients. This allowed us to understand how variation in among-species predation risk underlies different patterns of variation in community-level predation risk. To understand how habitat might relate to variation in prey susceptibility at each site, we calculated Pearson r correlation coefficients between mean eelgrass shoot density and percent cover of eelgrass, bare ground, and macroalgae and the community- and species-level variation in predation risk. Finally, to assess the role each individual species played in driving patterns of variation in predation risk at the community level (both aggregated and between the species), we calculated the standard deviation of the annual predation risk for each species at each site (i.e., the standard deviation of each colored line in Figure 2A) and correlated these values with the average percent cover of seagrass. This allowed us to see how the degree of seasonality of predation risk that prey items experienced changed across sites, and if this variation was related to habitat.

## Results

### Predation assays

Across all sites (n = 6) and time points (n = 16), we deployed and retrieved 5840 tethers over the course of one year. Variability in predation risk differed both among species and sites (Figure 2A). If prey items became unglued in the field as the result of some extreme environmental condition not present in these laboratory tanks, like high temperatures or strong currents, we would expect to see consistently high removal of all the species at all the sites during times of the year with peak water temperature (July – August; DuBois et al. 2022) or frequent storm events (February – March; Bromirski et al. 2003). However, there was wide variation in the predation risk observed during these periods, including some of the lowest values we observed across the whole year (Figure 2A). All this, in addition to the relative consistency we observed in predation risk when conducting assays on the same species at the same site at frequent intervals (Figure 2A), gives us confidence in assuming prey items were removed by consumption from predators, rather than some external environmental factor.

Across all sites, variation in among-species and community-level predation risk were negatively associated (r = −0.96, df = 4, p = 0.002; Figure 3A): sites with more consistent aggregate community predation risk had larger differences in among-species susceptibility on average across the year (Figure 3A). At sites with the largest differences in the risk experienced by the different species (BL, WP), there was very little variation in average predation risk of the whole community across the year (such as the scenarios depicted in Figure 1B or 1C). However, these sites also had the most variation in the distributions or “shape” of seasonal predation risk for each species (Figures 2B & 2C), indicating that the pattern is more like what is shown in Figure 1C than Figure 1B. On the other hand, at the site with the smallest differences among species at any point in time (NC), total community predation risk was highly variable across the year, reaching its lowest point in the spring and peaking in the fall (as depicted in Figure 1A).

**Figure 3.**
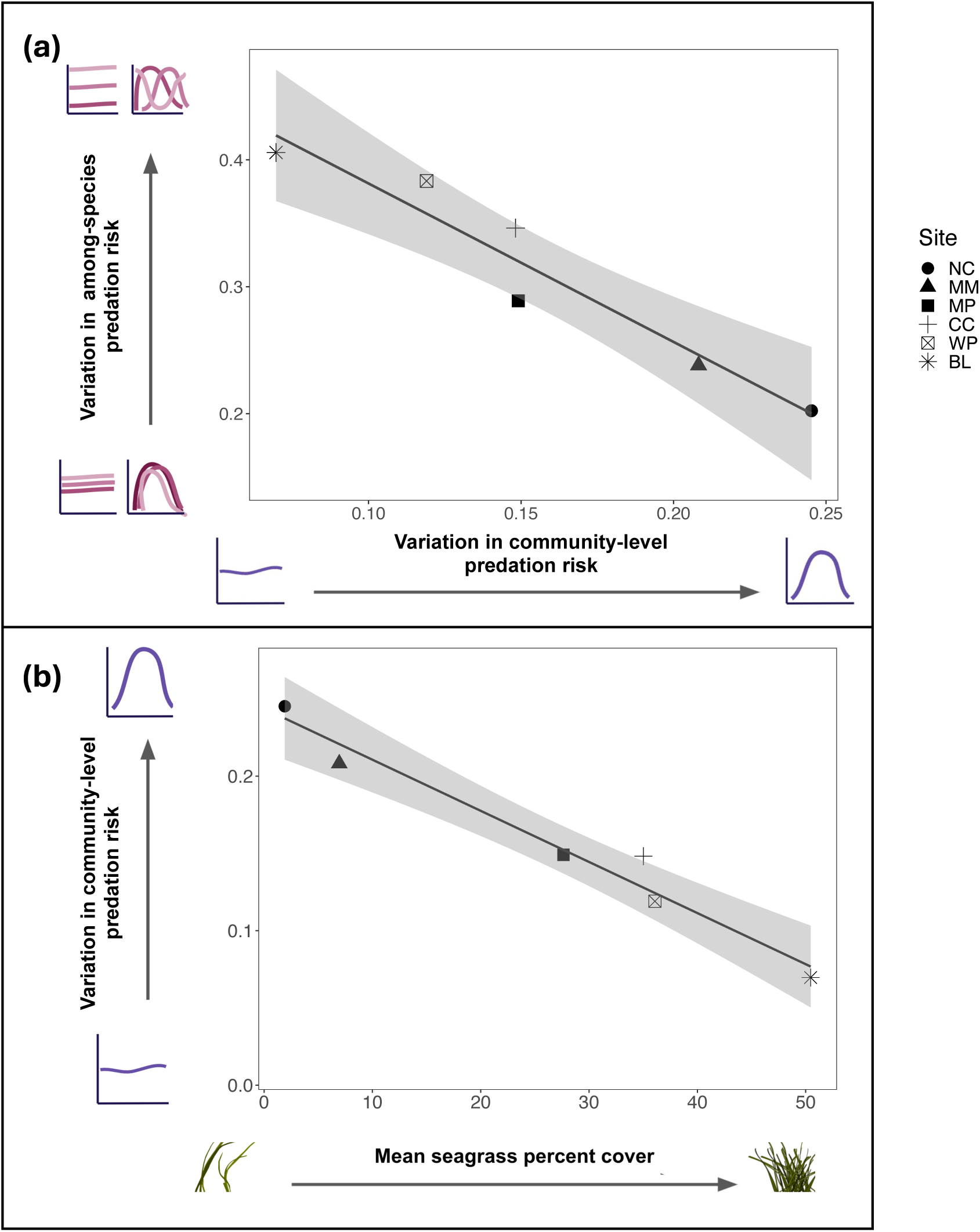
Increased stability in community-level predation risk is correlated with increased variation in among-species predation risk **(a)** and these are both correlated with increased seagrass cover **(b)**, a potential driver of this pattern. Each differently shaped dot corresponds to a different site. Shaded areas represent the 95% confidence interval around each line. Seagrass illustrations are by Claire E. Murphy.

When we ran the PCoA on our KS-distance matrix, only 12% of the total variation was explained by negative eigenvalues. We scaled the variation explained by the first two PCoA axes to be relative to the total variation explained by all the positive axes. Without this scaling, these first two axes explained 65% and 20% of the total variation explained, and after the scaling they represented 53% and 16% of the variation explained by all the positive axes.

### Habitat metrics

Sites varied in their average amount of seagrass cover across the year, ranging from 2% (annual range: 0% to 10% cover, at site NC; Appendix S1: Figure S2A) to 50% (annual range: 13% to 89% cover, at site BL; Appendix S1: Figure S2A). The average amount of cover at a site was negatively correlated with the variation in community-level predation risk (r = −0.963, df = 4, p = 0.002; Figure 3B) and positively correlated with the variation in species-specific predation risk (r = 0.975, df = 4, p < 0.001). Sites also varied in average shoot density, ranging from 1.15 (annual range: 0 to 3.8 at NC) to 34.8 (annual range: 13 to 100 at BL) shoots per 0.25 m^2^ (Appendix S1: Figure S2B). These values were also correlated with variation in predation risk both among species (r = 0.921, df = 4, p = 0.009) and at the community level (r = −0.946, df = 4, p = 0.004). Similarly, the average amount of bare ground, which can be used as a proxy for the total amount of structure-free space (with no eelgrass, algae, or large sessile invertebrates) ranged from 50.1% (at BL) to 94.0% (at NC) and increased with the community-level variation in predation risk (r = 0.884, df = 4, p = 0.019) but decreased as the among-species variation increased (r = −0.902, df = 4, p = 0.014). The mean percent cover of just macroalgae ranged from 0.067% (at BL) to 36.4% (at CC) and was not correlated with either metric of predation risk variability (among-species: r = 0.051, df = 4, p = 0.924; community-level: r = 0.114, df = 4, p = 0.829).

Considering each species separately across sites, annual variation in predation risk was correlated with habitat metrics for only two of the four species. The standard deviation of risk decreased with increasing percent eelgrass cover and shoot density for the isopod *P. resecata* and the caprellid *C. californica* (percent cover: r = −0.882, df = 4, p = 0.020; shoot density: r = - 0.797, df = 4, p = 0.056 and percent cover: r = −0.877, df = 4, p = 0.022; shoot density: r = - 0.917, df = 4, p = 0.010, respectively; Figure 4). In other words, how predation risk changed with the amount of habitat varied among species such that at the sites with least eelgrass, the predation risk of the isopods and caprellids becomes more temporally variable, which is more similar to the predation risk patterns of the ampithoids. This is also evident in the KS distance PCoA (Appendix S1: Figure S3), where the distributions of predation risks for the ampithoids overlap with the distributions of caprellids and isopods at the site with the least eelgrass, while the predation patterns for the isopods and caprellids are much more variable among sites, especially at those with highest eelgrass cover. All species converged on a similar seasonal pattern of predation risk at low habitat availability sites but were more differentiated at high habitat availability sites.

**Figure 4.**
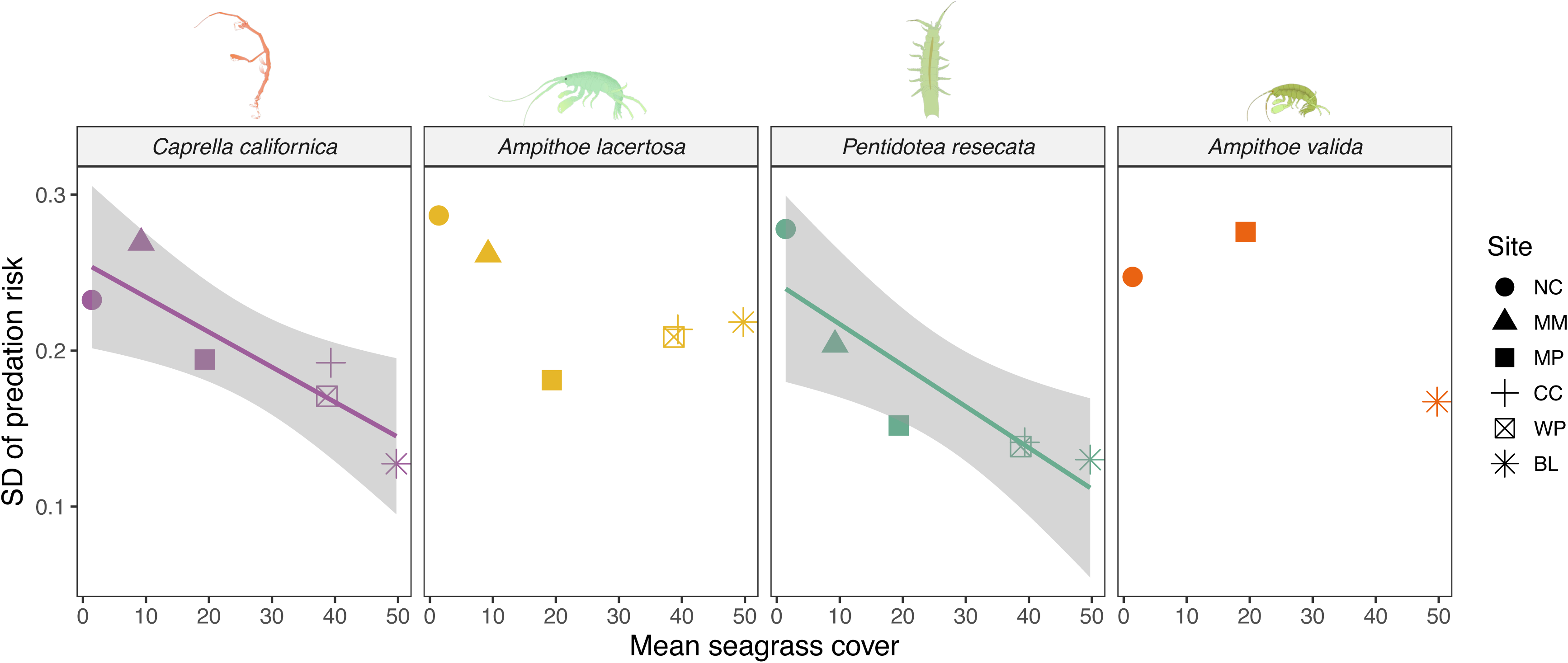
The standard deviation of the predation risk each species experiences at each site only decreases with increasing seagrass cover for the species with the most morphological defenses (*Pentidotea resecata* for both mean and standard deviation and *Caprella californica* for standard deviation). Trendlines with shaded areas representing the 95% confidence intervals are shown for the significant linear relationships. Crustacean illustrations are by Collin Gross.

## Discussion

We found that at sites where each prey species in the community had more distinct annual predation patterns, there were more stable levels of aggregate community predation risk across the year. We suggest a possible explanation for this pattern is that sites with high eelgrass cover facilitated the expression of species-specific predator avoidance strategies that depend on habitat. At sites with high habitat availability, each prey species differed in the timing of seasonal peaks and troughs in predation risk, creating complementarity which averaged to constant total (summed across all species) prey consumption through the year. At sites with low amounts of habitat, in contrast, all species had the same strongly seasonal pattern of predation risk, with a maximum in early fall and a minimum in winter. We suggest that these patterns may be a result of habitat amount influencing both the effectiveness of different antipredator strategies and the composition of seasonal predator communities and hypothesize that this altering of seasonal predation patterns could promote prey community diversity and stability.

The four species we used exhibit a range of predator avoidance strategies, which we speculate may contribute to the differences in predation risk patterns we observed at sites with varying amounts of habitat. At all sites, regardless of habitat amount, the two prey species with the most behavioral antipredator strategy (the ampithoids *A. lacertosa* and *A. valida*, which construct temporary silk tubes to hide in) were always consumed at the highest (Figure 2A) and most seasonally variable rates (Figure 4). In contrast, the species with the strongest morphological predator defenses (the isopod *P. resecata*, which has a large bulky body and camouflages well in seagrass; Lee and Gilchrist 1972), was consumed at relatively low, less seasonally variable rates at sites with high eelgrass cover (Figure 2A; Figure 4), resulting in higher among-species variation in predation risk at these sites. However, as the amount of habitat decreased, the strong defenses of this species seemed to be less effective, as it was eaten more frequently during the late summer, experiencing the same predation risk as the ampithoids (Figure 4) and reducing the overall among-species variation in predation risk at low eelgrass cover sites (Figure 3). The remaining prey species, the caprellid *C. californica*, was intermediate in how predation varied among sites with different eelgrass cover. Morphologically, caprellids are thin and thus relatively inconspicuous to visual predators when still, but behaviorally, when feeding or dispersing their jerky movement and large claw-like gnathopods may make them more easily detected by visual predators like fish (Virnstein et al. 1984, Caine 1989). This mix of traits translated to a similar pattern to what was seen with *P. resecata*, where *C. californica* was eaten at more seasonably variable rates as the amount of habitat decreased (Figure 4). However, *C. californica* consistently had a higher predation risk than *P. resecata* (Figure 2A), potentially due to weaker morphological defenses and their increased visibility.

When prey rely on distinct modes of predator avoidance, tethering may differentially interact with these strategies. For example, tethering could have reduced the effectiveness of behavioral avoidance strategies more than the morphological ones, potentially resulting in inflated estimates of predation risks. However, we do not believe this greatly influenced our results, as the hierarchy of predation risk among species is consistent with tether-free studies including both lab experiments and gut content analyses of prey selectivity (Nelson 1979, Stoner 1979, Best and Stachowicz 2012).

Seasonal changes in predation may also be influenced by an interaction between habitat and the abundance or behavior of predators. Though we did not measure the predator community abundance or composition here, eelgrass beds, like other complex, biogenic habitats, provide shelter for many small bodied fish in the summer months – both juveniles of more transient species and adults of localized resident ones (Lefcheck et al. 2019). This general increase in abundance of predator communities might explain the general shift in magnitude of predation risk between the winter and summer months. However, changing foraging behaviors across habitat contexts may explain the seemingly decreased effectiveness of predator avoidance strategies at the relatively barren sites. In addition to reduced effectiveness of defenses described above, generalist predators should become less selective in their prey choice as the costs of foraging increase (Prokopenko et al. 2023). Thus, mesopredators (such as the species that eat isopods and amphipods in our study) hunting in lower density habitats often become more generalist in their prey selection, regardless of available prey densities in the field (Holbrook and Schmitt 1988, Verdolin 2006), perhaps because they are more vulnerable to their own predators and thus spend less time selecting among prey. At the sites with more habitat, predators may be less vulnerable themselves and spend more time foraging (Mukherjee et al. 2009, Dhellemmes et al. 2021), which can promote selective feeding throughout the year. Overall, the high habitat cover throughout the year means that the site always has the potential to support a predator community that is selective of the prey they consume. This may have led to the temporally complementary patterns we observed in the predation risk at the high habitat sites (e.g., BL in Figure 2A, similar to Figure 1C), where each species was consumed at high rates at different points in the year.

The differences we observed in the seasonality of predation risk within the prey community may also influence prey abundance and co-occurrence. Across many marine predator types, increased habitat structural complexity has been shown to lead to a shift from type II to type III functional responses (Dunn and Hovel 2020), potentially due to the increased complexity facilitating low density refuges and prey-switching, common drivers of type III functional responses (Murdoch 1969). In denser and more structurally complex habitats, it is more important for predators to develop a specific search image to find their prey (Siddon and Witman 2004), which may be easier to do than for some of our prey species than others. Large dissimilarities in the costs of either finding or capturing different prey species also incentivizes predators to prey-switch (Prokopenko et al. 2023). These phenomena may help explain why we observed more variable seasonal predation patterns among prey species at sites with more habitat. Furthermore, all these drivers have been shown in other systems to lead to more stable prey populations (Oaten and Murdoch 1975). An intriguing hypothesis worthy of future testing is that increased habitat cover could mediate the stability of the prey community by amplifying the effects of different predator avoidance strategies and incentivizing prey-switching between higher density prey.

Overall, we found that sites with greater differences in their species-specific predation risk had more stable levels of community-level predation risk, seemingly driven by temporal complementarity. Furthermore, habitat cover was correlated with these differences. We suggest that increased habitat may mediate the importance of among-species differences in predator-avoidance strategies across the entire year, resulting variable timings of peak predation risk for different prey species and thus more stable levels of predation as a community-level process.

Different species are at an advantage across the year as they experience different periods of relief from predation, potentially mediating prey co-occurrence and acting as a mechanism for prey diversity maintenance. Our results suggest a series of hypotheses that, if tested, would improve our understanding of how interactions between refuge availability and prey traits, even without knowledge of the predator community, alter predation risk, leading to a better understanding of prey population dynamics across space and time.

## Supporting information

Video S1 Metadata

## Acknowledgements

Fieldwork for this project was conducted on the unceded traditional lands of the Coast Miwok people. We are grateful to many technicians and volunteers for their help with completing all the required lab and fieldwork, including Emma Deen, Sophie Allen, Lara Hsia, Cassidy Gordon, Collin Gross, Tracie Hayes, Adri Penix, Anna Lee, Lily McIntire, Keira Monuki, Naomi Murray, Sam Walkes, and Karolina Zabinski. Funding for this project came from a UC Davis Population Biology Grant to CEM and by National Science Foundation Awards OCE 18-29992 and 23-11578 to JJS.

## Author Contributions

CEM and JJS conceptualized the study. CEM collected the data, conducted the analyses, and drafted the manuscript with significant input from JJS.

