## Appendix S1 for "Seasonal differences in predation risk among seagrass epifauna species stabilize community-level predation over time"

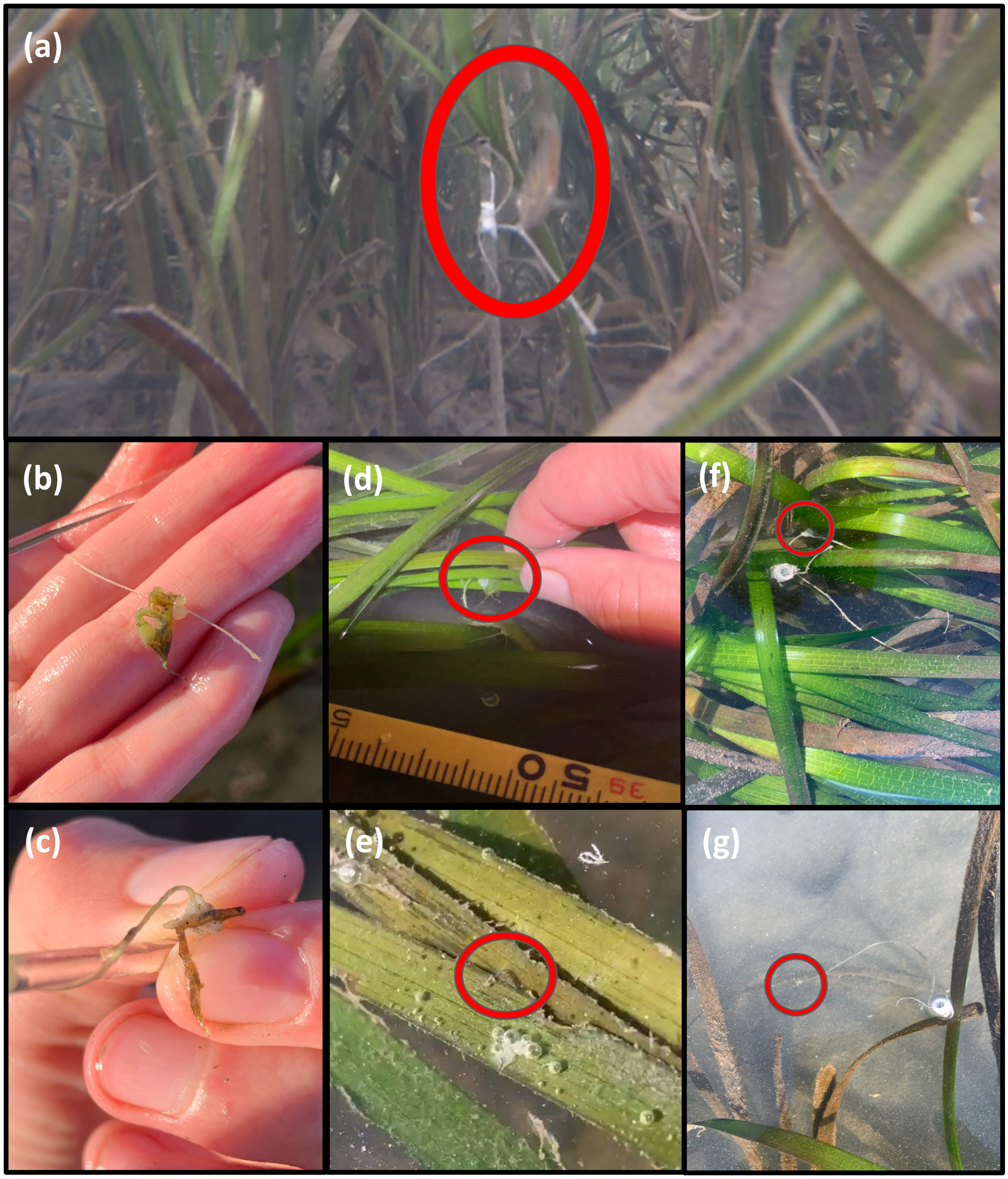


**Figure S1.** Various pictures from the field giving support to our conclusions that tethering assays measure the relative predation risks of our four species. These photos show shows a juvenile surf perch consuming a tethered *A. valida*, which is a still from Video S1 (**a**); partially consumed *A. lacertosa* and *C. californica* (**b** & **c**); a tethered ampithoid using blades of eelgrass to create a protective tube (**d**) in the same manner as an untethered ampithoid (**e**), both of which are circled in red; and tethered isopods (circled in red) finding eelgrass blades to cling to and hide on in both high and low habitat environments (**f** & **g**). Together these images show that tethered individuals can exhibit normal behaviors, and missing individuals at retrieval are likely removed due to predation. All photos by Claire E. Murphy.

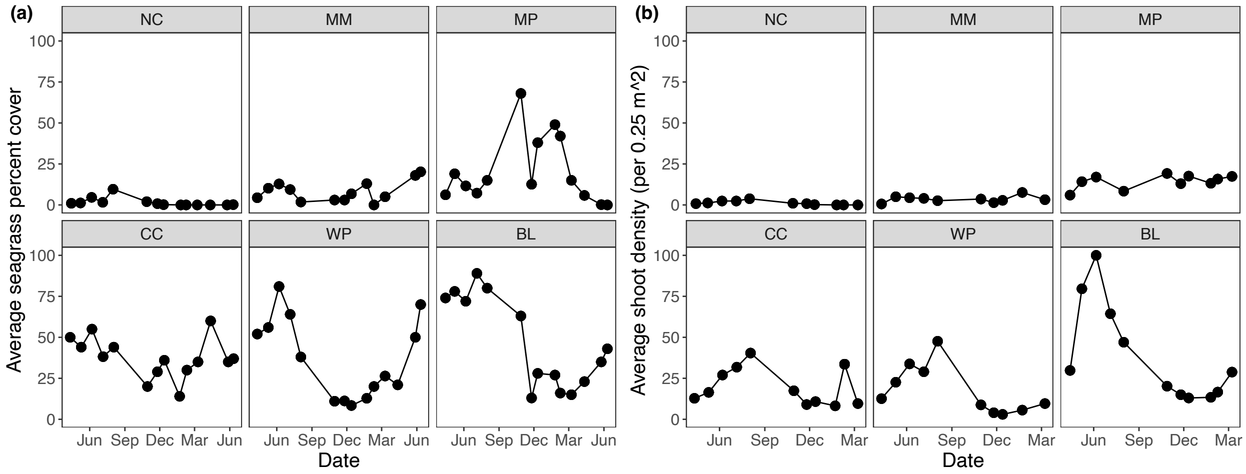


**Figure S2.** The average seagrass percent cover **(a)** and shoot density per 0.25 m$^2$ **(b)** across the year at each of the six sites, represented by the six boxes in each panel.

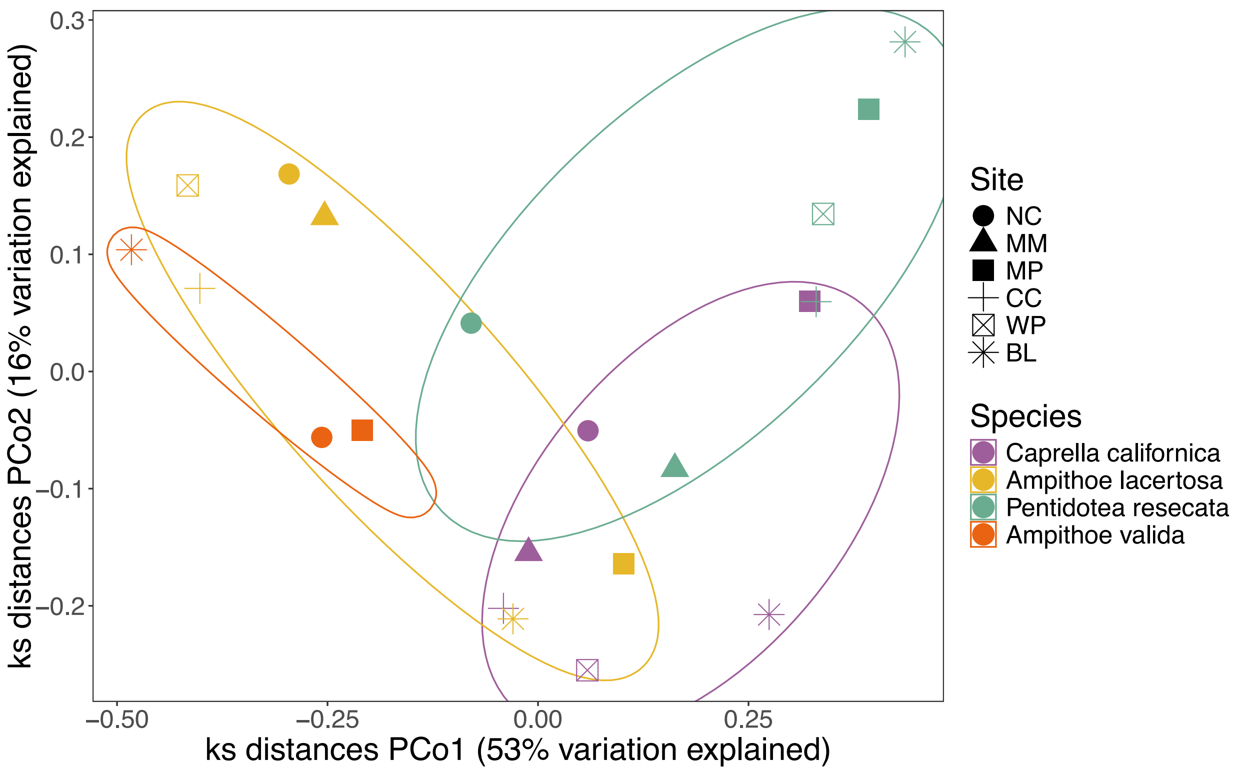


**Figure S3.** Each species by site predation pattern plotted in Kolmogorov-Smirnov (KS) multivariate space group more by species identity than by site. Each point represents a distribution of predation risk across the year (e.g., one of the colored lines in Fig. 2A) for each species (represented by colors) at every site (represented by shapes). The distance between two points correlates to how similar those two distributions are (e.g., the similarity of the shapes of two different colored lines in Fig. 2A).
