## Supplementary material for "Seasonal differences in predation risk among seagrass epifauna species stabilize community-level predation over time": Video S1 Metadata

**Video S1.** A shiner perch (*Cymatogaster aggregate*) can be seen eating a tethered *Ampithoe valida* before swimming off at a site with a high amount of eelgrass cover (Blake's Landing). Video was taken by Claire Murphy.
